# Janus Kinase 3 phosphorylation and the JAK/STAT pathway are positively modulated by follicle-stimulating hormone (FSH) in bovine granulosa cells

**DOI:** 10.1101/2022.02.05.479268

**Authors:** Amir Zareifard, Francis Beaudry, Kalidou Ndiaye

## Abstract

JAK3 is a member of the JAK family of tyrosine kinase proteins involved in cytokine receptor-mediated intracellular signal transduction through the JAK/STAT signaling pathway. JAK3 was shown as differentially expressed in granulosa cells (GC) of bovine pre-ovulatory follicles suggesting that JAK3 could modulate GC function and activation/inhibition of downstream targets. We used JANEX-1, a JAK3 inhibitor, and FSH treatments and analyzed proliferation markers, steroidogenic enzymes and phosphorylation of target proteins including STAT3, CDKN1B/p27^Kip1^ and MAPK8IP3/JIP3. GC were treated with or without FSH in the presence or not of JANEX-1. Expression of steroidogenic enzyme *CYP11A1*, but not *CYP19A1*, was upregulated in GC treated with FSH and both were significantly decreased when JAK3 was inhibited. Proliferation markers *CCND2* and *PCNA* were reduced in JANEX-1-treated GC and upregulated by FSH. Western blots analyses showed that JANEX-1 treatment reduced pSTAT3 amounts while JAK3 overexpression increased pSTAT3. Similarly, FSH treatment increased pSTAT3 even in JANEX-1-treated GC. UHPLC-MS/MS analyses revealed phosphorylation of specific amino acid residues within JAK3 as well as CDKN1B and MAPK8IP3 suggesting possible activation or inhibition post-FSH or JANEX-1 treatments. We show that FSH activates JAK3 in GC, which could phosphorylate target proteins and likely modulate other signaling pathways involving CDKN1B and MAPK8IP3.

## Introduction

The ovary is responsible for producing oocytes through a precise regulation by gonadotropins (follicle-stimulating hormone (FSH) and luteinizing hormone (LH)) and by steroid hormones (estrogen and progesterone) within the ovarian follicles^1–4^. In early stages of folliculogenesis, it is known that the ovarian follicles develop independently of gonadotropins while ovary-derived paracrine factors, such as cytokines and other growth factors play crucial roles in these early stages^5^. Although the beginning of folliculogenesis is gonadotropin-independent, receptors of gonadotropins are present in follicles prior to antrum formation advancing the importance of gonadotropins in early stages of follicular development^6^. As follicles progress into the antral stage, gonadotropins, especially FSH, become crucial for follicle survival and growth^6^. Granulosa and theca cells are two types of somatic cells that provide a suitable environment for the development of the oocyte within the follicle. These cells are responsive to gonadotropins and steroid hormones, which affect follicular development and ovulation for the release of a mature oocyte for subsequent fertilization^7–11^. Granulosa cells in particular are an important component of the follicle as they contribute to steroid hormone synthesis^12^, oocyte maturation^13^, and corpus luteum formation after ovulation^14^. The control of granulosa cells proliferation and function is therefore complex and depends on the precise regulation and activation or inhibition of specific genes and signaling pathways. FSH binds to its receptors on granulosa cells and initiates a series of molecular responses to regulate the expression of specific genes that are required for cell proliferation and growth as well as cumulus expansion and differentiation. While the initiation of follicular development is under the control of FSH, the release of the oocyte from the ovulatory follicle is under the influence of the LH surge (see ^3^ and ^4^ for reviews). However, the molecular mechanisms and underlying signaling pathways to these critical biological processes are still not fully investigated.

Previous *in vivo* gene expression analysis using bovine granulosa cells identified several candidate genes differentially regulated during different stages of follicular development^15–17^. Of interest, JAK3 was shown to be differentially expressed in granulosa cells of pre-ovulatory follicles and downregulated in ovulatory follicles by LH or hCG injection^17^ suggesting a potential role of JAK3 in regulating ovarian follicular development. JAK3 is a non-receptor tyrosine kinase that belongs to the JAK family proteins along with JAK1, JAK2 and TYK2^18^. JAK proteins have seven JAK homology (JH) domains namely the FERM, SH2, pseudokinase and kinase domains^19,20^. The FERM region at the N-terminus mediates attachment to cytokine receptors, while the SH2 domain provides a docking site for downstream proteins such as signal transducers and activator of transcription (STAT) proteins^21^. The kinase region bearing the activation loop at the carboxyl terminus and responsible for the tyrosine phosphorylation is located next to a pseudokinase region, which lacks any demonstrated kinase activity^22^. Within the JAK/STAT pathway and upon activation, JAK3 recruits and phosphorylates downstream effectors including STAT proteins, which leads to the regulation of target genes and modulation of cell proliferation, function and survival^23^. This well-defined pathway is yet to be fully investigated in reproductive cells including ovarian granulosa cells. Concurrently, additional JAK3 targets that can be affected by JAK3 activation in granulosa cells might include Cyclin-dependent kinase inhibitor 1B (CDKN1B also known as p27^Kip1^) and Mitogen-activated protein kinase 8 interacting protein 3 (MAPK8IP3) also known as JNK-interacting protein 3 (JIP3) previously identified as JAK3 binding partners^17^. CDKN1B binds to and prevents the activation of cyclin E-CDK2 or cyclin D-CDK4 complexes, and thus controls the cell cycle progression at G1^24^. CDKN1B is therefore often referred to as a cell cycle inhibitor protein since its major function is to stop or slow down the cell division cycle when it is not phosphorylated. As for MAPK8IP3, it was shown to interact with and regulate the activity of numerous protein kinases of the c-Jun N-terminal kinase (JNK) signaling pathway and thus functions as a scaffold protein in signal transduction^25^. The JNK pathway is one of the major signaling cascades of the mitogen-activated protein kinase (MAPK) signaling pathway and regulates a number of cellular processes including proliferation, embryonic development and apoptosis^26^.

Overall, the available data regarding JAK3 regulation in reproductive cells point to a crucial role in controlling granulosa cells activity and proliferation during follicular development prior to the LH surge and subsequent ovulation and luteinization processes. Previous studies have shown the importance of JAK signaling in the formation of primordial follicles as well as germline cyst breakdown as inhibition of JAK3 decreased pregranulosa cell formation through the downregulation of NOTCH2 signaling^27^. Other studies have shown stronger abundance of STAT3 phosphorylation in granulosa cells of small follicles after follicle deviation as related to granulosa cells death and follicular atresia^28^. However, within the selected follicle, the exact role of JAK3 in the regulation of granulosa cells activity and phosphorylation of target proteins still remains poorly defined. We hypothesized that JAK3 could modulate granulosa cells proliferation and steroidogenic activity as well as activation or inhibition of downstream targets. It is well established that phosphorylation on serine, threonine and tyrosine residues is an extremely important modulator of protein function as these modifications can be critical in the activation or inactivation of proteins of interest^29^. Therefore, the present study was conducted to unveil the function and mechanism of action of JAK3 in granulosa cells during ovarian follicular development. The study was conducted using inhibition and overexpression strategies as well as stimulation of granulosa cells to elucidate the regulation and phosphorylation sites within target proteins. We show that JAK3 influences granulosa cells proliferation, steroidogenesis activity and function through phosphorylation of target proteins, activation of the JAK/STAT pathway and likely modulation of other signaling pathways involving CDKN1B and MAP8IP3 in response to stimulations such as FSH. This study provides evidence that significantly deepen our molecular understanding of JAK3 activity and role in the regulation of a proper ovarian follicular development and mammalian folliculogenesis.

## Methods

### Chemicals and Products

Rabbit antibodies against STAT3 (Cat. # ab32500) and phospho STAT3 (Cat. # ab32143) were from Abcam. Anti-β-actin antibodies (Cat. # SC-47778) used as loading reference was from Santa Cruz Biotechnology. Follicle-stimulating hormone (FSH; Cat. # F4021-10UG) were purchased from Sigma while the pharmacological inhibitor JANEX-1 was purchased from Santa Cruz Biotechnology (SC-205354, LOT # B1011). Complete protease inhibitors were purchased from Roche Diagnostics (Laval, QC, Canada). M-PER Mammalian Protein Extraction Reagent (Cat. # 78503) was obtained from Thermo Fisher.

### Experimental Animal Models

Animal Ethics Committee of the Faculty of Veterinary Medicine of the University of Montreal reviewed and approved all the experimental protocol as the cows were cared for in accordance with the Canadian Council on Animal Care guidelines^30^. The expression and regulation of all JAK family members was analyzed during different stages of follicular development and ovulation using an *in vivo* model previously described^15,17^. Granulosa cells were collected from a dominant follicle group (DF, n = 4) and an hCG-induced ovulatory follicle group (OF, n = 4) obtained from ovaries of synchronized cows^31^. DF samples were obtained from the ovaries bearing the dominant follicle on day 5 of the estrous cycle (day 0 = day of estrus) as previously reported^15^. Follicles were dissected in order to isolate granulosa cells (GC). The OF samples were obtained following an injection of 25 mg of PGF_2α_ (Lutalyse) on day 7 of the estrous cycle to induce luteolysis, thereby promoting the development of the dominant follicle of the first follicular wave into a preovulatory follicle. An ovulatory dose of hCG (3000 IU, iv; APL, Ayerst Lab, Montréal, QC) was injected 36 h after the induction of luteolysis, and the ovary bearing the hCG-induced OF was collected 24 hours post-hCG injection for GC isolation. Samples were stored at −80°C until further analyses. Furthermore, GC were collected from 2 to 4mm small antral follicles (SF) obtained from slaughterhouse ovaries, and a total of three pools of 20 SF were prepared. Corpora lutea (CL; n=3) were collected at day 5 of the estrous cycle from the same cows used for DF sampling and stored at −80°C. These samples are referred to as *in vivo* samples and were used to validate JAK family members regulation during follicular development.

### Functional Analyses

For *in vitro* experiments, GC were aspirated from small to medium size follicles (< 8 mm in diameter) of slaughterhouse ovaries and put into culture in DMEM/F12 medium. GC were cultured in 12-well plates (n=4 independent experiments with duplicate wells for each treatment) for RNA and protein extraction. Cells were seeded at a density of 0.1×10^6^ cells/well. The complete DMEM/F12 culture medium was supplemented with L-glutamine (2 mM), sodium bicarbonate (0.084%), bovine serum albumin (BSA; 0.1%), HEPES (20 mM), sodium selenite (4 ng/mL), transferrin (5 μg/mL), insulin (10 ng/mL), non-essential amino acids (1 mM), androstenedione (100 nM), penicillin (100 IU) and streptomycin (0.1 mg/mL) as previously published ^32,33^. For FSH treatment, cells were incubated in complete culture medium supplemented with insulin (10 ng/mL) for 2 days, then in a medium containing FSH (1 ng/mL) and insulin (10ng/mL) for 4 days. In JAK3 inhibition experiments, granulosa cells were cultured and treated with 100 μM of JANEX-1. This concentration was based on previously published data^17^. JANEX-1 treatment was either followed by FSH treatment (1 ng/mL) or not. Cells for all experiments were incubated at 37°C in a humidified 5% CO_2_ atmosphere for a total of 6 days with media changed every 48 hours. Effects of FSH and JANEX-1 treatments on GC were assessed by analyzing expression of *JAK3* by RT-qPCR and JAK3 downstream targets STAT3, CDKN1B and MAPK8IP3 phosphorylation status by western blot or HPLC/MS-MS. Cell proliferation markers *cyclin D2 (CCND2)* along with *proliferating cell nuclear antigen (PCNA)* were also evaluated by RT-qPCR analyses. To further analyze GC steroidogenic activity, steroidogenic genes *CYP19A1* (P_450_ aromatase) and *CYP11A1* (P_450_ cholesterol side-chain cleavage) were measured by RT-qPCR.

For overexpression experiments, the JAK3 open reading frame was amplified by PCR using the Advantage 2 polymerase mix (Takara Bio) with JAK3-ORF primers (Table 1) amplifying the entire 3372 bp open reading frame. The JAK3 PCR fragment was purified and cloned into the pQE-Tri System His-Strep2 vector (Qiagen). The final pQE-JAK3 construct was used to transfect GC using the Xfect transfection kit (Takara Bio) according to the manufacturer’s protocol. The effects of JAK3 overexpression were assessed by measuring phosphorylation levels of STAT3.

**Table 1.**
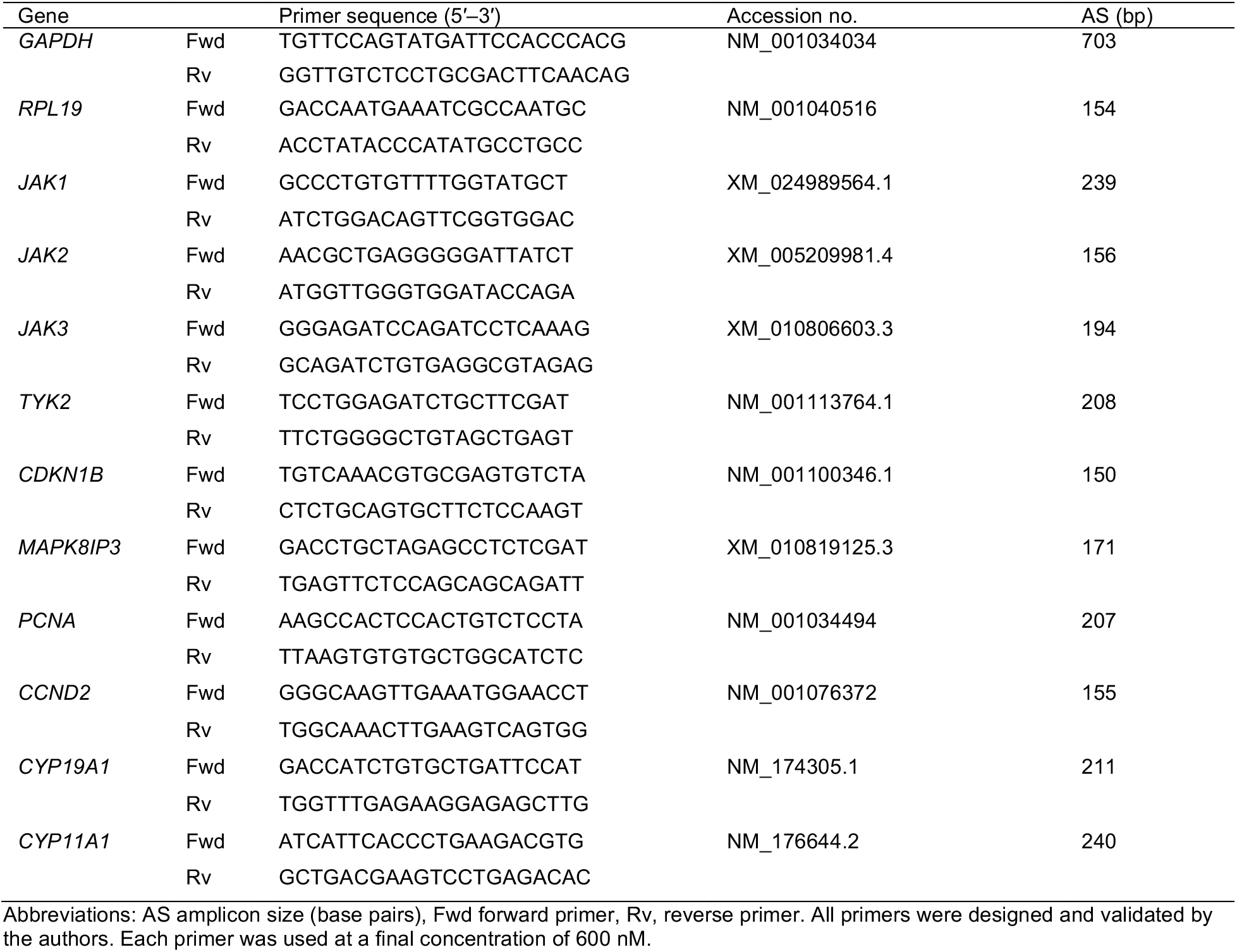
Primers used in the expression analyses of Bos taurus target genes by PCR and RT-qPCR

### Proteomic Analysis: Cell extracts and UHPLC-MS/MS

Protein pellets from control cells, FSH- and JANEX1-treated cells were dissolved in 100 μL of 50 mM TRIS-HCl buffer (pH 8), and the solution was mixed with a Disruptor Genie at maximum speed (2,800 rpm) for 15 min and sonicated to improve the protein dissolution yield. Proteins samples were denatured by heating at 120°C for 10 min using a heated reaction block and allowed to cool for 15 min. Proteins were reduced with 20 mM dithiothreitol (DTT), and the reaction was performed at 90°C for 15 min. Then, proteins were alkylated with 40 mM iodoacetamide (IAA) protected from light at room temperature for 30 min. Then, 5 μg of proteomic-grade trypsin was added, and the reaction was performed at 37°C for 24 h. Protein digestion was quenched by adding 10 μL of a 1% trifluoroacetic acid (TFA) solution. Samples were centrifuged at 12,000 *g* for 10 min, and 100 μL of the supernatant was transferred into injection vials for analysis using a Thermo Scientific Vanquish FLEX UHPLC system (San Jose, CA, USA). Chromatography was performed using gradient elution along with a Thermo Biobasic C18 microbore column (150 × 1 mm) with a particle size of 5 μm. The initial mobile phase conditions consisted of acetonitrile and water (both fortified with 0.1% formic acid) at a ratio of 5:95. From 0 to 3 min, the ratio was maintained at 5:95. From 3 to 123 min, a linear gradient was applied up to a ratio of 40:60, which was maintained for 3 min. The mobile phase composition ratio was then reverted to the initial conditions, and the column was allowed to re-equilibrate for 30 min. The flow rate was fixed at 50 μL/min, and 5 μL of each sample was injected. A Thermo Scientific Q Exactive Plus Orbitrap Mass Spectrometer (San Jose, CA, USA) was interfaced with the UHPLC system using a pneumatic-assisted heated electrospray ion source. Nitrogen was used as the sheath and auxiliary gases, which were set at 15 and 5 arbitrary units, respectively. The auxiliary gas was heated to 300°C. The heated ESI probe was set to 4000 V, and the ion transfer tube temperature was set to 300°C. Mass spectrometry (MS) detection was performed in positive ion mode operating in TOP-10 data dependent acquisition (DDA) mode. A DDA cycle entailed one MS^1^ survey scan (m/z 400-1500) acquired at 70,000 resolution (FWHM) and precursor ions meeting the user-defined criteria for charge state (i.e., z = 2, 3 or 4), monoisotopic precursor intensity was selected for MS^2^ acquisition (dynamic acquisition of MS^2^-based TOP-10 most intense ions with a minimum 2×10^4^ intensity threshold). Precursor ions were isolated using the quadrupole (1.5 Da isolation width), activated by HCD (28 NCE) and fragment ions were detected in the ORBITRAP at a resolution of 17,500 (FWHM). Data were processed using Thermo Proteome Discoverer (version 2.4) in conjunction with SEQUEST using default settings unless otherwise specified. The identification of peptides and proteins with SEQUEST was performed based on the reference protein sequences proteome extracted from UniProt (JAK3: E1BEL4_BOVIN, CDKN1B: A6QLS3_BOVIN, MAPK8IP3: A0A3Q1LQI8_BOVIN) as FASTA sequences. Parameters were set as follows: MS^1^ tolerance of 10 ppm; MS^2^ mass tolerance of 0.02 Da for Orbitrap detection; enzyme specificity was set as trypsin with two missed cleavages allowed; carbamidomethylation of cysteine was set as a fixed modification; and oxidation of methionine as well as phosphorylation of serine, threonine and tyrosine were set as a variable modification. The minimum peptide length was set to six amino acids. Data were further analyzed with a target-decoy database to filter incorrect peptide and protein identifications. For protein or peptide quantification and comparative analysis, we used the peak integration feature of Proteome Discoverer 2.4 software.

### mRNA Expression Analysis

Expression and regulation of *JAK3* and other JAK family members (*JAK1, JAK2* and *TYK2*) mRNA were analyzed by RT-qPCR during follicular development using *in vivo* samples and specific primers (Table 1). Total RNA was extracted from bovine GC collected from follicles at different developmental stages (SF, DF, OF) and CL as described above and previously published ^34^. Total RNA were also extracted from *in vitro* samples of cultured GC and reverse transcription reactions were performed using the SMART PCR cDNA synthesis technology (Takara Bio.) according to the manufacturer’s procedure and as previously published ^35^. *In vitro* samples were used to analyze the expression of JAK3 binding partners (*CDKN1B* and *MAPK8IP3*), proliferation markers (*PCNA* and *CCND2*) as well as steroidogenesis markers (*CYP19A1* and *CYP11A1*). RT-qPCR experiments were performed using SsoAdvanced Universal SYBR Green Supermix (Bio-Rad) following the manufacturer’s protocol. RT-qPCR data were analyzed using the Livak method (2^-ΔΔct^)^36^ with *RPL19* used as reference gene^37^.

### Cell Extracts and Immunoblotting Analysis

Cultured GCs were collected and homogenized with the M-PER mammalian protein extraction reagent (Thermo Fisher) supplemented with complete protease inhibitors. Total proteins from the different treatment groups were extracted and cell debris were removed by centrifugation (14,000 × *g* for 5 min at 4 °C) and the supernatants were collected and stored at −80 °C until further analyses. Total protein concentrations were determined using the Bradford method^38^ (Bio-Rad Protein Assay, Bio-Rad Lab, Mississauga, ON, Canada) and immunoblotting was performed as described previously^39^. Protein samples (50 μg) were resolved by one-dimensional denaturing 12 % Novex Trisglycine gels (Invitrogen, Burlington, ON, Canada) and transferred onto polyvinylidene difluoride membranes (PVDF; GE Healthcare Life Sciences). Membranes were incubated with specific first antibodies against STAT3 at a concentration of 0.069 μg/mL and phospho-STAT3 (pSTAT3) at a concentration 1 μg/mL to verify the effects of JAK3 inhibition or activation on the JAK/STAT pathway. Membranes were also incubated with anti-beta actin (ACTB) as reference protein. Immunoreactive proteins were visualized by incubation with appropriate horseradish peroxidase-linked secondary antibodies (1:10000 dilution) followed by incubation with the chemiluminescence system (Thermo Scientific) according to the manufacturer’s protocol and revelation was done using the ChemiDoc XRS+ system (Bio-Rad).

### Experimental Design and Statistical Rationale

Data are presented as mean ± SEM from three or more independent experiments, unless otherwise specified in the text. Values for *JAK3* and other target genes mRNA were normalized to the reference gene *RPL19*. Homogeneity of variance between groups was verified by O’Brien and Brown-Forsythe tests. Corrected values of gene specific mRNA levels were compared between follicular or CL groups using one-way analysis of variance (ANOVA). When ANOVA indicated a significant difference (p ≤ 0.05), the Tukey-Kramer test was used for multiple comparison of individual means among SF, DF, OF, and CL and for *in vitro* experiments, whereas the Dunnett test (p ≤ 0.05) was used to compare different samples from UHPLC-MS/MS. For Western blot analyses, cells were grown to appropriate confluence and treated as described. Protein bands were quantified using ImageJ software (NIH), corrected with beta-actin as loading reference and compared using ANOVA. For proteomics analyses, data sets were further analyzed with a target-decoy database to filter incorrect peptide and protein identifications. For protein or peptide quantification and comparative analysis, we used the peak integration feature of Proteome Discoverer 2.4 software. Statistical analyses were performed using GraphPad PRISM version 9 for macOS.

## Results

### JAK family members are differentially regulated during follicular development

Previously, we reported that JAK3 was differentially expressed in GC of preovulatory follicles and downregulated in ovulatory follicles *in vivo* by luteinizing hormone (LH) or following human Chorionic Gonadotropin (hCG) injection^15,17^. To generate an overall analysis of the regulation of JAK family members in bovine granulosa cells, total RNA was extracted from small follicles (SF), dominant follicles at day 5 of the estrous cycle (DF), ovulatory follicles obtained 24h post-hCG injection (OF) and corpus luteum (CL) at day 5 of the estrous cycle in order to analyze *JAK1, JAK2, JAK3* and *TYK2* expression. RT-qPCR analyses showed that JAK family members are differently regulated during different stages of follicular development. Gene expression analyses revealed the strongest expression of *JAK3* in SF and DF and its significant downregulation in OF and CL (Fig. 1a; p ≤ 0.001), confirming previously reported data from our laboratory^17^. In addition, *JAK1* was induced by hCG in OF as compared to SF, DF and CL (Fig. 1b; p ≤ 0.001). *JAK1* expression was also stronger in DF as compared to SF (p ≤ 0.05). *JAK2* expression was substantially present in the CL as compared to small, dominant and ovulatory follicles (Fig. 1c; p ≤ 0.0001) while among the different groups of follicles, *JAK2* expression was stronger in OF as compared to SF and DF (Fig. 1c; p ≤ 0.05). As for *TYK2* expression, it was stronger in DF and OF compared to the SF and CL samples (Fig. 1d; p ≤ 0.05). These results suggest that JAK members might play different functions in the ovary and modulate different downstream targets in GC and therefore might not present functional redundancy. Specifically, JAK3 might be involved in GC proliferation and follicular development while other JAK members might play significant roles in the ovulation or luteinization processes or maintenance of a functional corpus luteum.

**Figure 1.**
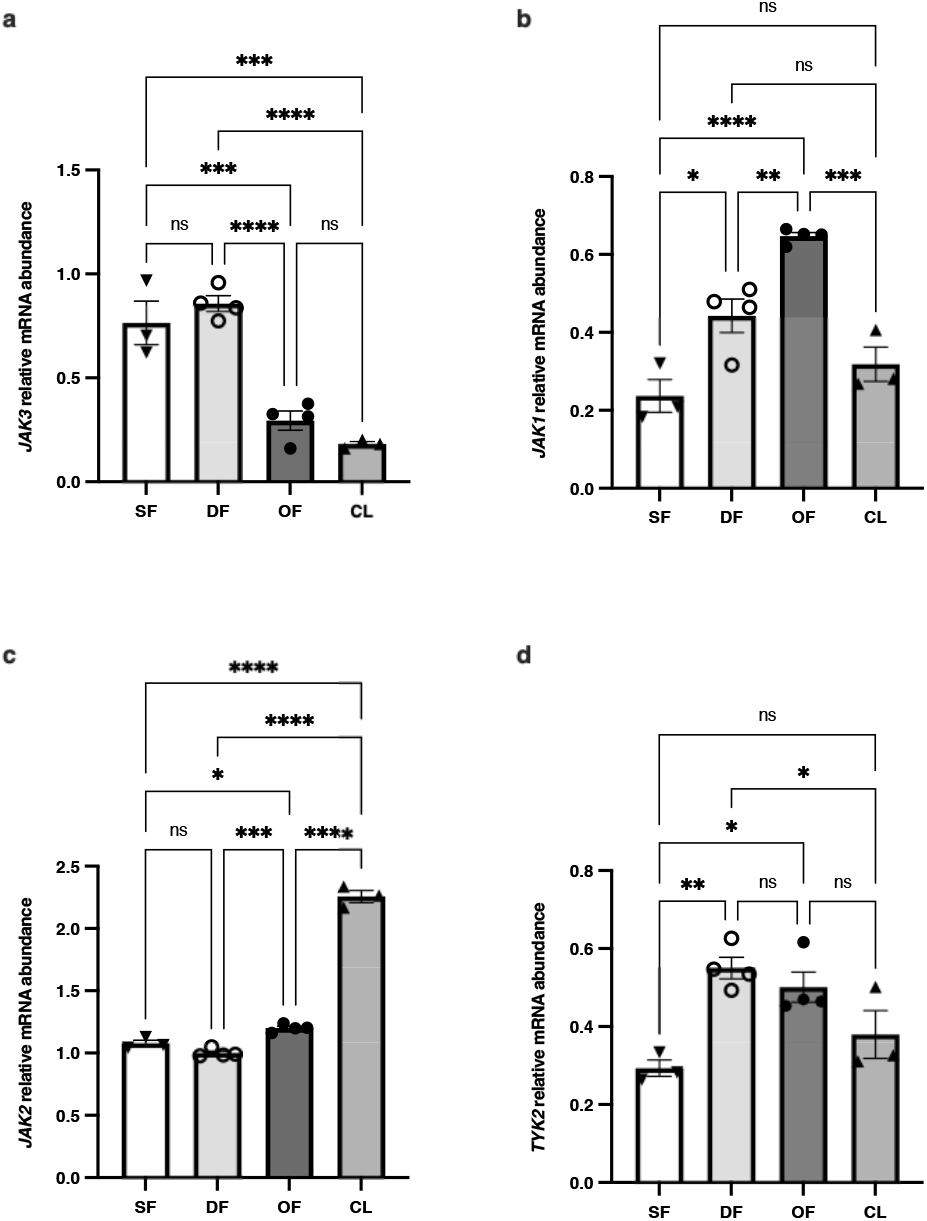
Regulation of *JAK* family members during follicular development. mRNA expression and regulation of JAK family members were analyzed in GC of bovine follicles at different stages using RT-qPCR. Total RNA was extracted from GC of small follicles (SF; N=3), dominant follicles (DF; N=4), ovulatory follicles (OF; N=4), and corpus luteum at day 5 (CL; N=4) as described in materials and methods and target genes were analyzed (*JAK1, JAK2, JAK3, TYK2*) relative to *RPL19* as reference gene. *JAK3* mRNA expression was strongest in DF and SF and was significantly decreased in OF and CL (p ≤ 0.05) while other JAK members were differentially expressed in different stages of follicular development and CL. This result suggests a different regulation of JAK family in granulosa cells at different stages of follicular development and CL. RT-qPCR data are presented as gene abundance using the 2^-ΔΔCt^ method. *, p ≤ 0.05, **, p ≤ 0.01; ***, p ≤ 0.001; ****, p ≤ 0.0001 (ANOVA, Tukey-Kramer multiple comparison); ns, not significant.

### JAK3 inhibition reduces expression of proliferation markers while FSH stimulates proliferation markers in cultured granulosa cells

JAK3 function in GC was analyzed using JANEX-1 (JNX), a specific JAK3 pharmacological inhibitor. RT-qPCR analyses were used to measure proliferation markers *cyclin D2 (CNND2)*, a cell cycle activator and *proliferating cell nuclear antigen (PCNA)* expression. The results showed that inhibition of JAK3 led to significant decrease in GC proliferation demonstrated by a decrease in both *CCND2* (Fig. 2a; p ≤ 0.0001) and *PCNA* (Fig. 2b; p ≤ 0.0001) expression as compared to the control group. FSH significantly increased both *CNND2* (Fig. 2a; p ≤ 0.01) and *PCNA* (Fig. 2b; p ≤ 0.05) expression as compared to the control and following JNX treatment. *PCNA* was used to assess and confirm GC proliferation since it is expressed in the nuclei of cells during the DNA synthesis phase of the cell cycle, which could be an indication of cell proliferation in GC. Moreover, expression of *CCND2* was also analysed using the same *in vitro* samples as another proliferation marker, which functions as a regulator of cyclin-dependent kinases required for cell cycle G1/S transition. Negative impacts of JAK3 inhibition on expression of *CCND2* and *PCNA* in JNX-treated GC suggest that JAK3 plays a central role in the regulation of ovarian granulosa cells proliferation and function. We also showed that FSH stimulation of cell proliferation following JNX inhibitory effects resulted in an increase in proliferation markers expression in GC (Fig. 2a and b) likely through activation of the JAK/STAT pathway. While we couldn’t detect a stimulatory effect of FSH on *JAK3* expression at the mRNA level, FSH clearly tended to increase *JAK3* expression following JNX treatment, likely as a consequence of FSH stimulation of cell proliferation following JNX inhibitory effects resulting in an increase in *JAK3* expression in GC (Fig. 2c; p ≤ 0.01).

**Figure 2.**
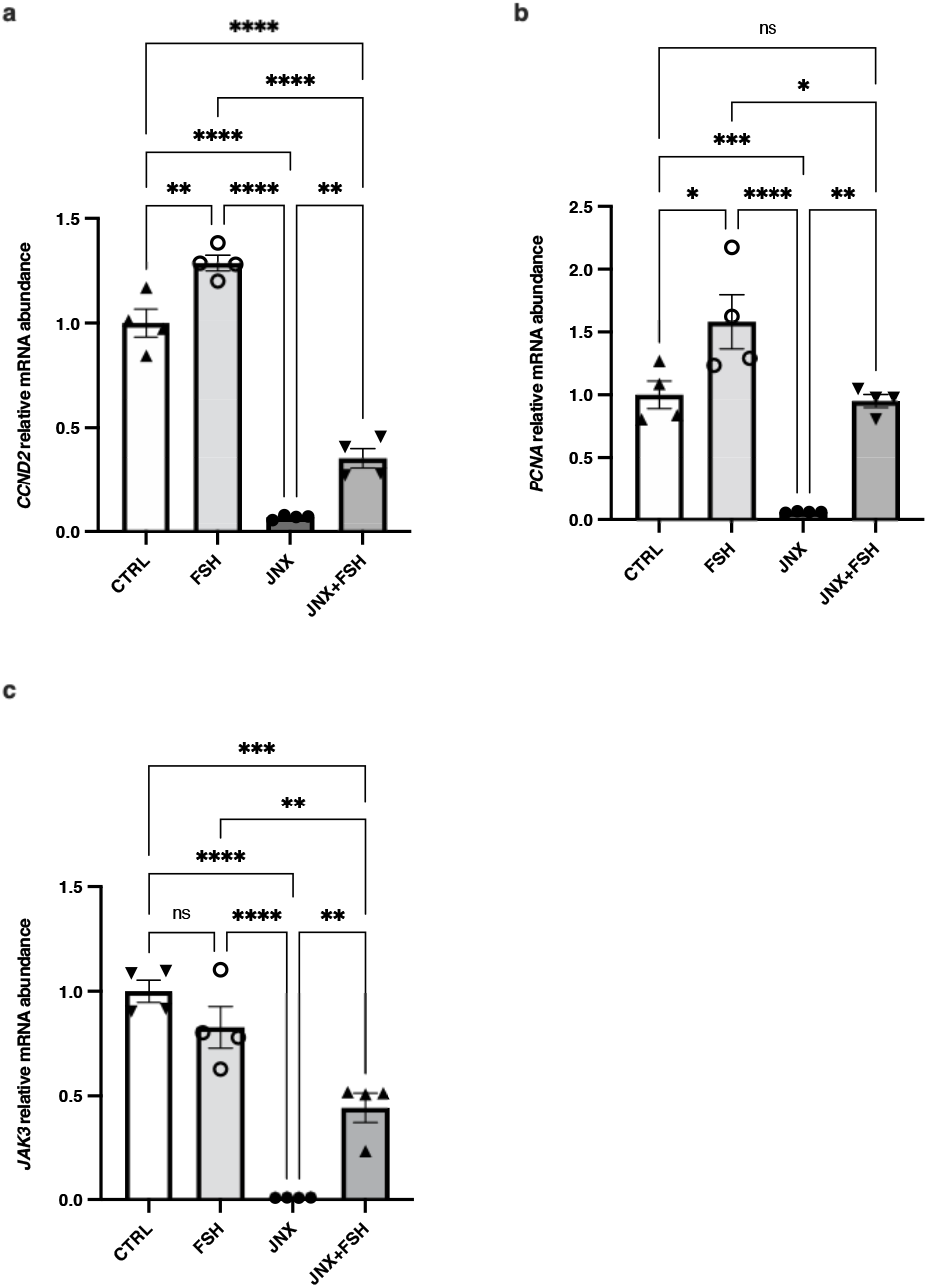
Regulation of *JAK3* mRNA expression *in vitro*. Total RNA was extracted from cultured GC treated with or without FSH or JNX and analyzed by RT-PCR for the expression of proliferation markers Cyclin-D2 (*CCND2*) and Proliferating cell nuclear antigen (*PCNA*) as well as *JAK3* relative to *RPL19* as reference gene. FSH stimulated both *CCND2* (a) and *PCNA* (b) mRNA expression levels in GC while JANEX-1 significantly decreased both of these proliferation markers after 24 h. In addition, addition of FSH following JNX treatment significantly increased expression of both *CCND2* and *PCNA* as compared to JNX. RT-qPCR analysis showed that *JAK3* mRNA expression in GC (c) significantly decreased after JNX treatment as compared to the control and FSH-treated cells. FSH treatment alone did not stimulate *JAK3* expression as compared to control group; however, FSH treatment significantly increased *JAK3* expression following JNX treatment. CTRL, control; FSH, Follicle-stimulating hormone; JNX, JANEX-1 RT-qPCR data are presented as gene abundance using the 2^-ΔΔCt^ method. *, p ≤ 0.05, **, p ≤ 0.01; ***, p ≤ 0.001; ****, p ≤ 0.0001 (ANOVA, Tukey-Kramer multiple comparison); ns, not significant.

### JAK3 inhibition modulates granulosa cells steroidogenic activity

To verify the effects of JAK3 inhibition on the steroidogenic activity of GC *in vitro,* expression of *CYP11A1* and *CYP19A1* enzymes were analysed as steroidogenesis markers and important regulators of the development and function of mammalian ovaries. RT-qPCR analyses showed that expression of both *CYP11A1* and *CYP19A1* was significantly decreased when JAK3 was inhibited with JNX as compared to the control (Fig. 3a and b; p ≤ 0.0001). Conversely, FSH stimulated the steroidogenic activity of treated GC and significantly increased the expression level of *CYP11A1* (Fig 3a; p ≤ 0.001), but not *CYP19A1* (Fig 3b), as compared to the control. Although *CYP19A1* expression was not significantly increased by FSH treatment alone, the results demonstrated that *CYP19A1* expression was significantly increased in cells treated with FSH following JNX treatment (Fig. 3b; P ≤ 0.05), a result that mirrors JAK3 regulation in Fig. 2c. These data confirm a positive effect of FSH on GC and suggest a potential role of JAK3 in regulating the expression of steroidogenic enzymes and its involvement in follicular development and differentiation as its inhibition is associated with negative effects on GC steroidogenic activity.

**Figure 3.**
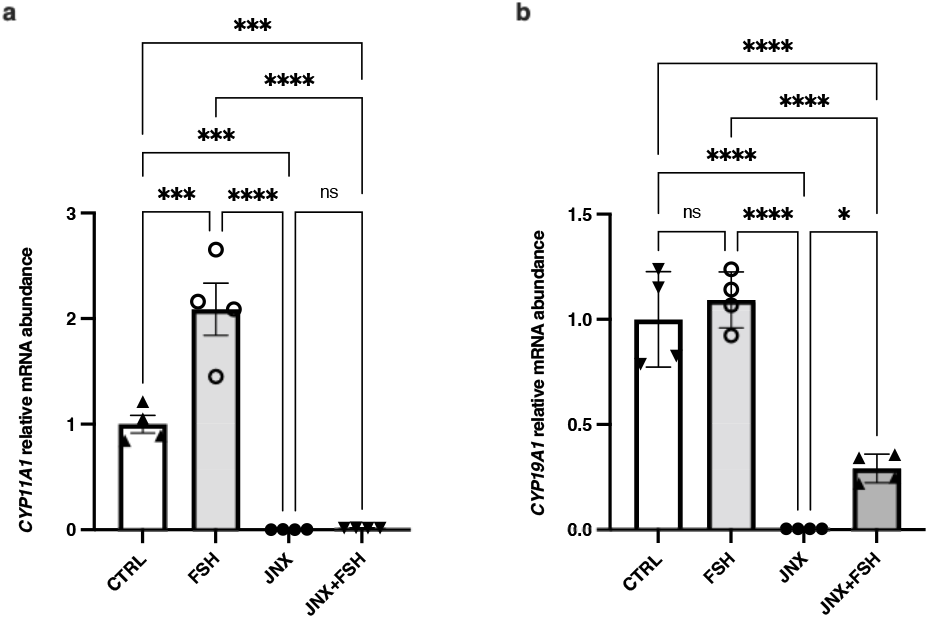
Effect of JAK3 inhibition on steroidogenesis markers (*CYP11A1* and *CYP19A1*) in granulosa cells. Total RNA was extracted from cultured GC treated with or without FSH or JANEX-1 and analyzed by RT-PCR for the expression of steroidogenic enzymes cytochrome P450 family 11 subfamily A member 1 (*CYP11A1*) and cytochrome *P450* family 19 subfamily A member 1 *(CYP19A1)* normalized against the house keeping gene *RPL19* as reference. The results show that FSH significantly upregulated *CYP11A1* in GC (a); however, *CYP19A1* (b) did change following FSH treatment. In addition, JAK3 inhibition with JNX resulted in a significant decrease of both *CYP11A1* and *CYP19A1.* FSH treatment following JANEX-1 significantly increased *CYP19A1* expression but not *CYP11A1.* CTRL, control; FSH, Follicle-stimulating hormone; JNX, JANEX-1. RT-qPCR data are presented as gene abundance relative to RPL19 using the 2^-ΔΔCt^ method. *, p ≤ 0.05, **, p ≤ 0.01; ***, p ≤ 0.001; ****, p ≤ 0.0001 (ANOVA, Tukey-Kramer multiple comparison); ns, not significant.

### JAK3 inhibition negatively affects STAT3 phosphorylation while JAK3 overexpression and FSH positively affect STAT3 phosphorylation in granulosa cells

To further analyse the possible regulatory influence of JAK3 on downstream effectors in GC, total extracted protein of *in vitro* samples was subjected to immunoblotting analyses. Relative abundance of STAT3 phosphorylation (pSTAT3) was quantified and presented as an indication of either STAT3 activation or inhibition within the JAK/STAT signaling pathway following overexpression or inhibition of JAK3 in GC using the pQE system or JNX treatment, respectively. Phosphorylation of STAT3 was also analysed and quantified following FSH treatment for 4 hours. The results showed that at 8h post JNX treatment, a significant decrease in pSTAT3 was observed as compared to the control while overexpression of JAK3 significantly increased pSTAT3 levels as compared to JNX group and back to the level of the control group (Fig. 4a; p ≤ 0.05). However, JNX treatment for 24 hours significantly decreased pSTAT3, whether in JAK3-overexpresed cells or not (Figure 4a; p ≤ 0.05, red bar vs black bar) meaning that JAK3 overexpression was not enough to reverse JNX effects after 24 hours of treatment as compared to 8 hours. In addition, phosphorylation of STAT3 was also analysed and quantified following FSH treatment alone, or in combination with JNX treatment. FSH administration for 4 hours increased pSTAT3 to the same level as JAK3 overexpression compared to JNX-treated GC (Figure 4b; p ≤ 0.0001). Treatment with FSH also significantly increased pSTAT3 amounts as compared to JNX-treated cells even after 24h as compared to JAK3 overexpression alone post-JNX treatment (Fig. 4b; p ≤ 0.0001, blue bar vs. black bar). This result suggests a specific effect on STAT3 phosphorylation levels via JAK3 modulation. These data support a role of FSH in positively modulating JAK/STAT signaling pathway by increasing pSTAT3 amounts in treated cells. The results also confirm that STAT3 phosphorylation levels are affected by JAK3 manipulation and suggest a central role of JAK3 in regulating the transmission of initiated signals through a functional JAK/STAT signaling pathway in GC, which would contribute to GC proliferation and follicular development. Overall, JNX negative effects on the JAK/STAT pathway were less drastic in the presence of FSH although JNX tended to decrease pSTAT3 in FSH-treated/JAK3-overexpressed granulosa cells (Fig. 4b; p = 0.0535).

**Figure 4.**
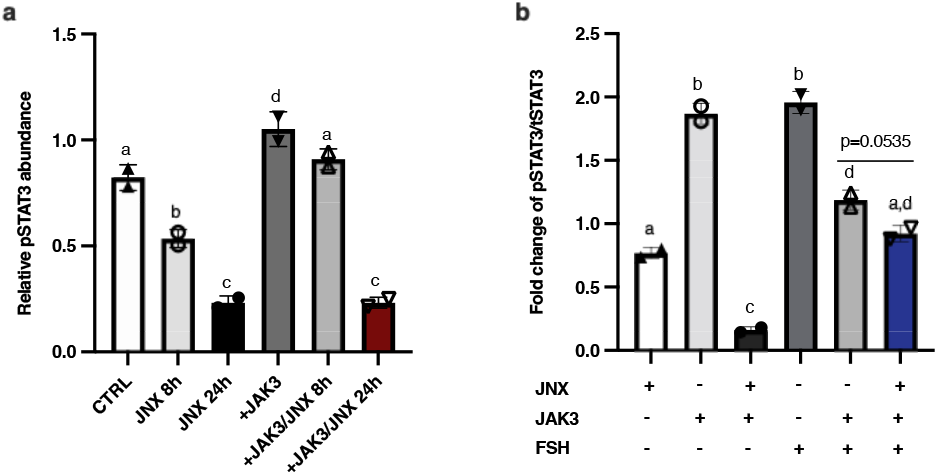
Effects of JAK3 inhibition or overexpression and FSH treatment on STAT3 phosphorylation in granulosa cells. **a)** To analyse the effects of JAK3 manipulations on downstream effectors in GC, Western blot analyses were performed and revealed phosphorylation status of STAT3 following different treatments. JAK3 was overexpressed in GC using the pQE system (+JAK3) with or without JNX treatment for 8 or 24 hours to inhibit JAK3 phosphorylation. Detected bands and their relative abundances were quantified and phosphorylation status of STAT3 (pSTAT3) was analysed. A significant increase in pSTAT3 amounts was observed when JAK3 was overexpressed. JNX treatment for 8 hours resulted in a significant decrease in pSTAT3. Overexpression of *JAK3* following 8 hours JNX treatment restored pSTAT3 amounts to the control levels. However, overexpression of *JAK3* was not able to restore or increase pSTAT3 within GC after 24 hours of JNX treatment, which significantly decreased pSTAT3 as compared to the control group and JAK3-overexpressed cells (red box). **b)** Further experiments allowed to evaluate the effects of FSH administration on the phosphorylation status of STAT3 in cultured GC. Fold changes of pSTAT3 over to total STAT3 were compared following treatments with JNX (24h), FSH (4h) or JAK3 overexpression. JNX significantly reduced phosphorylation of STAT3 after 24h of treatment, while overexpression of *JAK3* without JNX treatment significantly increased the amount of pSTAT3 within transfected GC similar to the result in A. Addition of FSH significantly increased pSTAT3 compared to the JNX group. Moreover, combination of JAK3 overexpression with FSH treatment increased pSTAT3 but not to the same levels as either FSH alone or JAK3 overexpression alone. Addition of FSH in the presence of JNX weakened the negative effect of JNX on pSTAT3 as compared to JAK3 overexpression in the presence of JNX (blue bar). CTRL, control; FSH, Follicle-stimulating hormone; JNX, JANEX-1; +JAK3, JAK3 overexpression. Different letters denote significant differences among samples (ANOVA, Tukey-Kramer multiple comparison).

### JAK3 inhibition differentially impacts binding partners CDKN1B & MAPK8IP3

UHPLC-MS/MS analyses revealed relative total peptide abundances as well as modifications at the amino acid levels including phosphorylation status of JAK3 and binding partners CDKN1B/p27^Kip1^ and MAPK8IP3/JIP3. Total abundances of these target proteins were analysed in GC treated with FSH or JNX as compared to a control group. Analysis and quantification of JAK3 protein revealed a substantial increase in total abundance following FSH treatment as compared to control and JNX treatment (Figure 5a and b), which suggests an increase in JAK3 phosphorylation and activation of JAK3 signaling pathway in FSH-treated GC. In contrast, JNX treatment for 24h notably decreased the abundance of JAK3 protein in treated granulosa cells as compared to the control (Fig. 5a; p = 0.094) and FSH treatment (Fig. 5a; p ≤ 0.05). Furthermore, peptides from JAK3 protein were recovered with different modifications including phosphorylation in specific amino acid residues. Quantification analysis of these recovered peptides revealed their abundances following treatments with FSH and JNX. Although not all JAK3 phosphorylated peptides showed significant changes in their abundances following FSH or JNX treatments when compared to the control or to each other, these phosphorylated fragments could indicate phosphorylation status of JAK3 leading to its activation and contribution to the subsequent molecular mechanisms. More specifically, the abundance of peptide #10.2 (amino acids 391-403 within JAK3 sequence) was not significantly affected following FSH or JNX treatment but exhibited phosphorylation on S^394^ and S^398^ residues and a relative decrease following JNX and (Fig. 5c). Similarly, peptide #23 (amino acids 871-887 within JAK3 sequence) exhibited phosphorylation on S^880^ and Y^886^ residues but there were no changes in its abundance following FSH or JNX treatment (Fig. 5d). Conversely, other phosphorylated JAK3 fragments were showing significant changes in their abundances and phosphorylation status following FSH treatment as compared to the control and JNX treatment, suggesting a more stringent regulation of these JAK3 regions. Of interest, peptide #20 (amino acids 735-748 within JAK3 sequence) was phosphorylated at Y^738^ and was significantly less abundant following JNX treatment as compared to FSH and control (Fig. 5e; p ≤ 0.05). Similarly, peptide 24 (amino acids 877-895 within JAK3 sequence) phosphorylated at S^4^ tended to be less abundant in JNX-treated cells as compared to FSH-treated cells and control (Figure 5f).

**Figure 5.**
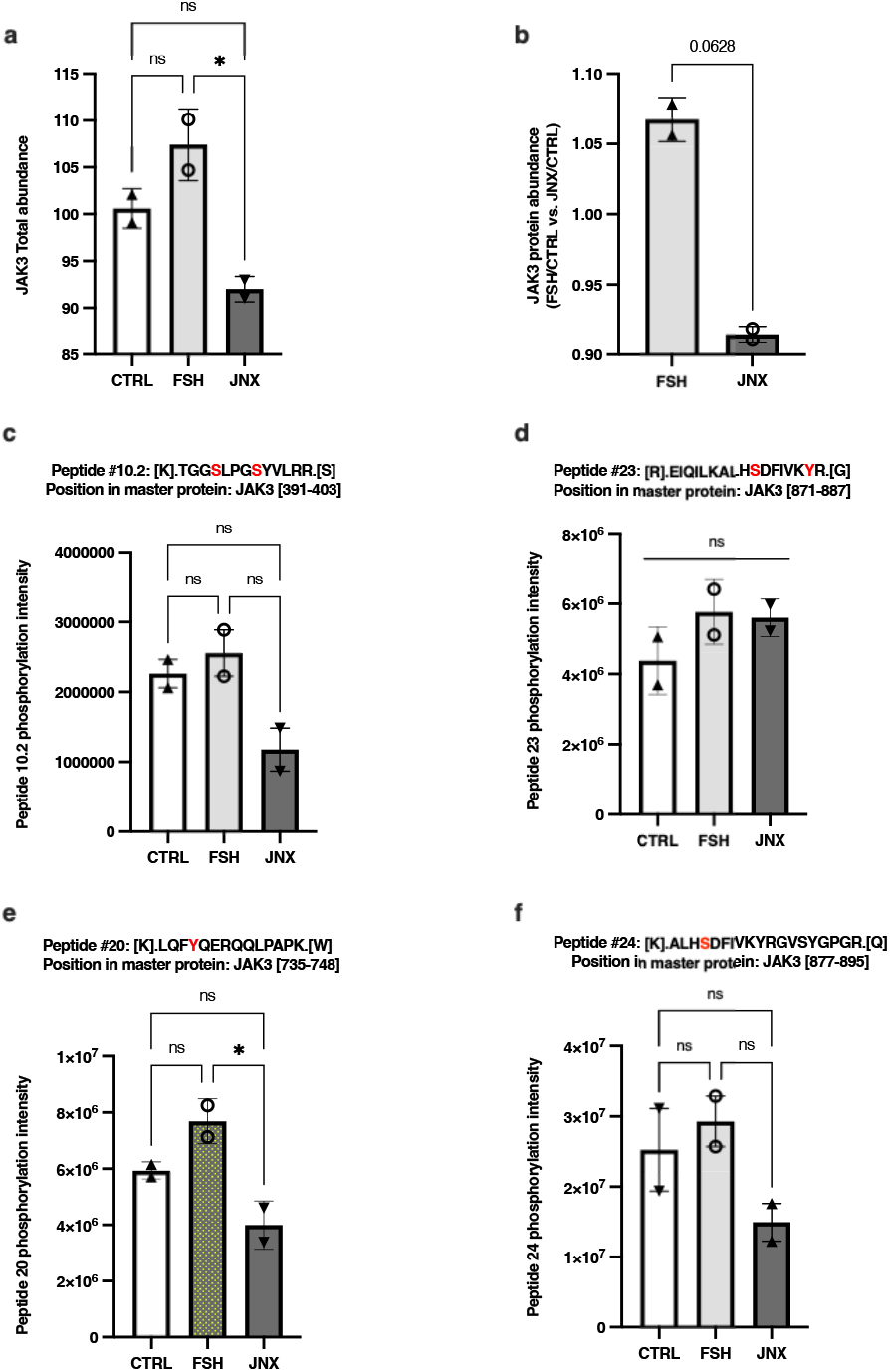
Effects of FSH and JANEX-1 treatments on JAK3 total abundance and peptides modifications in granulosa cells. HPLC/MS-MS was used to analyze and quantify total abundances of various recovered peptides for JAK3 in GC samples following treatments with FSH or JNX. **a)** JAK3 total abundance slightly increased following FSH treatment but not significantly while JNX treatment significantly decreased JAK3 abundance. **b)** Ratio of fold change in JAK3 abundance in FSH-treated cells over the control and in JNX-treated cells over the control are compared and showed JAK3 as relatively more abundant following FSH treatment (p = 0.0628). **c-f)** Peptides abundance and phosphorylation at specific amino acid residues. The abundances of various JAK3 phosphorylated peptides were quantified and evaluated following FSH or JNX treatments. Peptide numbers and positions in the JAK3 protein sequence are shown. Phosphorylated residues are shown in red. CTRL, control; FSH, Follicle-stimulating hormone; JNX, JANEX-1. *, p ≤ 0.05 (ANOVA, Tukey-Kramer multiple comparison); ns, not significant.

In addition, total protein abundances of CDKN1B and MAPK8IP3 were analysed in the same sets of *in vitro* samples than JAK3. The results revealed that total abundance of CDKN1B, which negatively affects cell cycle progression, is considerably decreased with FSH treatment (Fig. 6a and b; p = 0.075) and significantly increased post-JNX treatment (Fig. 6a and b; p ≤ 0.05), while MAPK8IP3 total abundance did not change significantly with FSH treatment but decreased significantly in JNX-treated cells as compared to control and FSH-treated cells (Fig. 7a and b; p ≤ 0.05). Further analyses of recovered peptides of CDKN1B and MAPK8IP3 revealed modifications at specific amino acid residues related to treatments with FSH or JNX. Abundance of peptide #1 (amino acids 6-15 within CDKN1B sequence) was increased following JNX treatment as compared to control and FSH treatment (Fig. 6c; p ≤ 0.05) but no phosphorylation was noted in any residue of this peptide suggesting a possible active CDKN1B protein in these JAK3-inhibited cells. However, peptide #7 (amino acids 190-198 within CDKN1B sequence) was phosphorylated at T^9^ and was significantly increased in FSH-treated cells as compared to JNX-treated cells (Fig. 6d; p ≤ 0.05) suggesting an inactivation of CDKN1B via this specific residue in the presence of FSH, which correlates with FSH positive effect on JAK3 activation and cell proliferation. As for peptides recovered for MAPK8IP3, peptides #9 (amino acids 201-219 within MAPK8IP3 sequence) and #34 (amino acids 595-605 within MAPK8IP3 sequence) exhibited phosphorylation modifications, respectively at residues S^17^ (peptide #9) and S^1^ and T/S^4/8^ (peptide #34) and both peptides were significantly decreased following JNX treatment (Fig. 7c and d; p ≤ 0.05) but did not change following FSH as compared to control (Fig. 7c). These data suggest that FSH stimulates JAK3 phosphorylation, which leads to the activation of the JAK/STAT signaling pathway through increased STAT3 phosphorylation as one of the downstream effectors, while likely inducing MAPK8IP3 activation and CDKN1B inactivation through phosphorylation of specific amino acid residues leading to cell cycle progression and granulosa cells proliferation.

**Figure 6.**
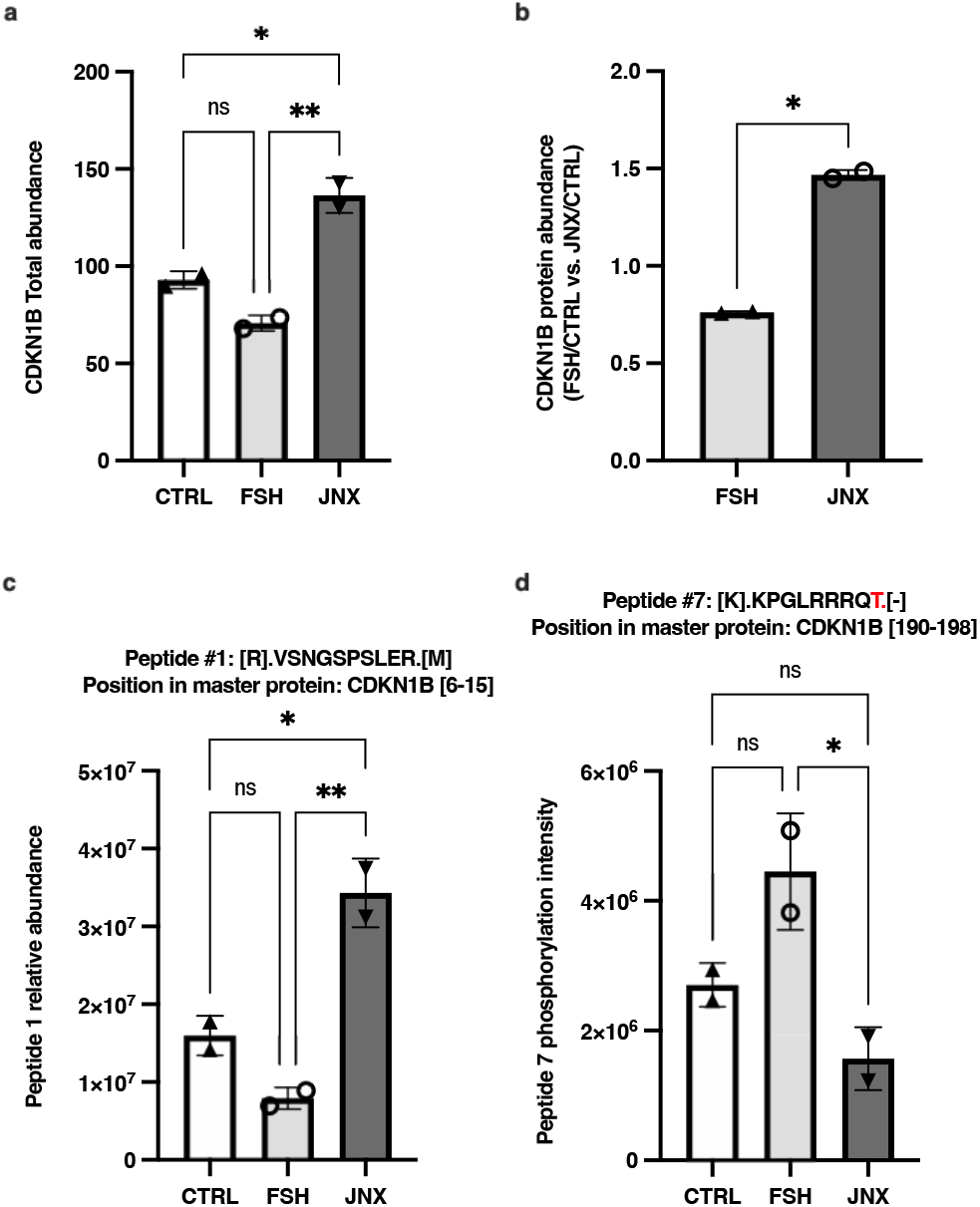
Effects of FSH and JANEX-1 treatments on CDKN1B total abundance and peptides modifications in granulosa cells. HPLC/MS-MS was used to analyze and quantify total abundances of various recovered peptides for CDKN1B in GC samples following treatments with FSH or JNX. **a)** CDKN1B total abundance tended to decrease following FSH treatment (P=0.075) while CDKN1B was significantly more abundant in JNX-treated cells. **b)** Ratio of fold change in CDKN1B abundance in FSH-treated cells over the control and in JNX-treated cells over the control are compared and showed CDKN1B more abundant following JNX treatment. **c-d)** Peptides abundance and phosphorylation at specific amino acid residues. The abundances of two CDKN1B phosphorylated peptides were quantified and evaluated following FSH or JNX treatments. Peptide numbers and positions in the CDKN1B protein sequence are shown. Phosphorylated residues are shown in red. CTRL, control; FSH, Follicle-stimulating hormone; JNX, JANEX-1. *, p ≤ 0.05; **, p ≤ 0.01 (ANOVA, Tukey-Kramer multiple comparison); ns, not significant.

**Figure 7.**
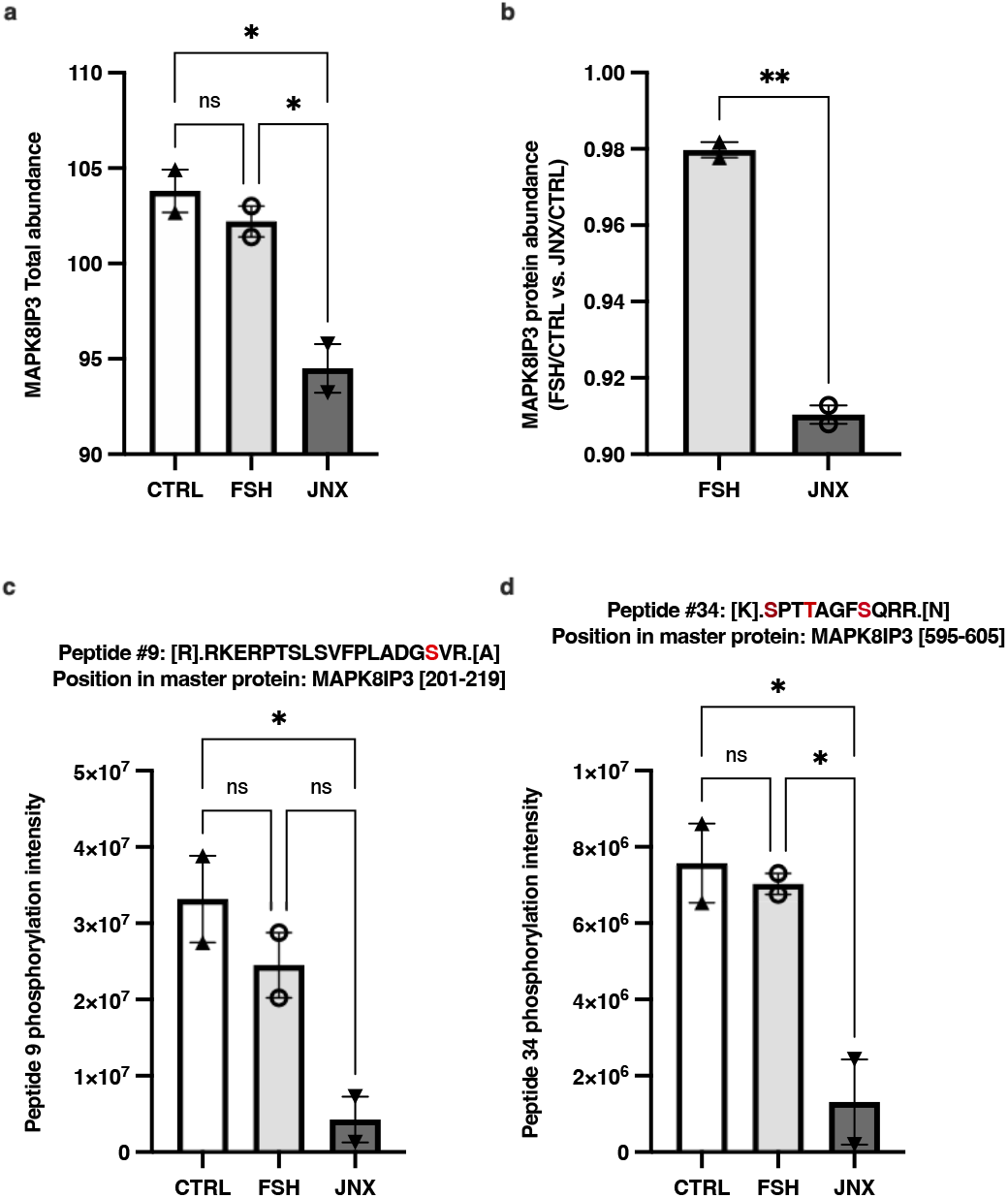
Effects of FSH and JANEX-1 treatments on MAPK8IP3 total abundance and peptides modifications in granulosa cells. HPLC/MS-MS was used to analyze and quantify total abundances of various recovered peptides for MAPK8IP3 in GC samples following treatments with FSH or JNX. **a)** MAPK8IP3 total abundance did not change following FSH treatment but JNX treatment significantly decreased MAPK8IP3 abundance. **b)** Ratio of fold change in MAPK8IP3 abundance in FSH-treated cells over the control and in JNX-treated cells over the control are compared and showed MAPK8IP3 as significantly more abundant following FSH treatment. **c-d)** Peptides abundance and phosphorylation at specific amino acid residues. The abundances of two MAPK8IP3 phosphorylated peptides were quantified and evaluated following FSH or JNX treatments. Peptide numbers and positions in the MAPK8IP3 protein sequence are shown. Phosphorylated residues are shown in red. CTRL, control; FSH, Follicle-stimulating hormone; JNX, JANEX-1. *, p ≤ 0.05; **, p ≤ 0.01 (ANOVA, Tukey-Kramer multiple comparison); ns, not significant.

### Conserved modifications for JAK3 and CDKN1B suggest activation and inactivation status in granulosa cells

Amino acid sequences of JAK3 and CDKN1B from bovine and human species were aligned using protein alignment tools (Clustal: Multiple Sequence Alignment) in order to analyse the similarities in phosphorylation sites and conserved areas between these two species. The analysis of recovered peptides from bovine samples revealed modifications at specific locations that were associated with JAK3 activation and CDKN1B inactivation in human^40–42^ and are conserved in the bovine species. As shown in the section above, these modifications were observed in this study following treatments with FSH or JNX and could be indicative of the JAK/STAT pathway activation or inactivation of CDKN1B in GC. Amino acid sequences alignment for JAK3 showed residues previously shown to be phosphorylated in human, which were retrieved in this study using bovine samples (Figure 8a; peptide 29 and related peptides). Modifications include phosphorylation of Tyrosine residues at 980-981 (Y^980-981^) in JAK3 following FSH treatment, which could be an indication of JAK3 activation. As shown in Figure 5, peptides in different regions of JAK3 were detected and were differentially regulated following the various treatments suggesting a potential phosphorylation pattern in different regions of JAK3 as an indication of JAK3 activity modulation.

**Figure 8.**
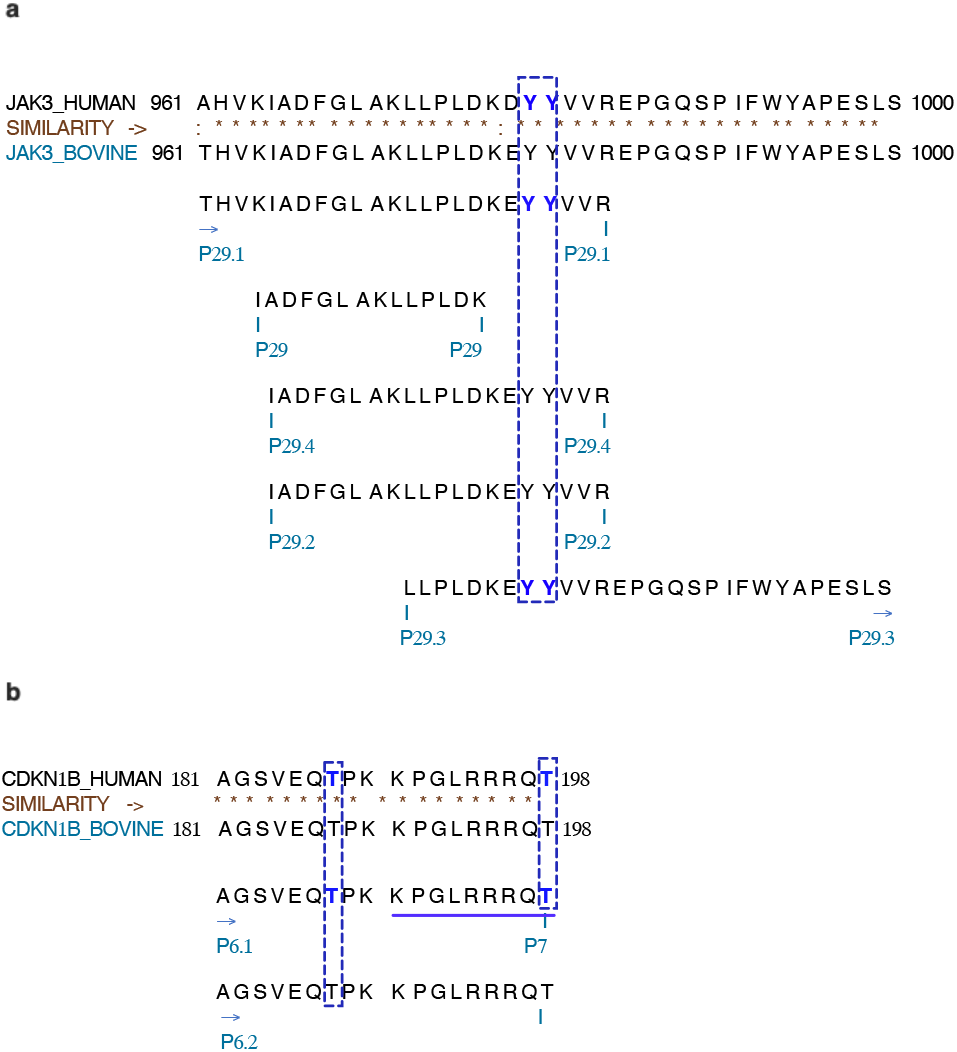
Recovered peptides for JAK3 and CDKN1B from HPLC/MS/MS analysis. **a)** Peptides sequences of JAK3 from human and bovine species were aligned to evaluate similarities and conserved modifications associated with JAK3 activation. Peptides positions within the JAK3 sequence are indicated and phosphorylated residues are shown in blue. In particular, phosphorylation of the two Tyrosine (Y) residues at locations 980-981 (peptide #29.1 in the figure) of JAK3 amino acid sequence in human were retrieved in this study. Similarly, different peptides with residues known to be phosphorylated in the human sequence were also found in this study meaning that these amino acid residues were also phosphorylated in bovine granulosa cells following JAK3 activation with FSH. **b)** For CDKN1B, the two highlighted Threonine (T) residues at locations 187 and 198 within peptide #7 are an indication of phosphorylation status of CDKN1B and are related to CDKN1B inactivation. Phosphorylated of this peptide was more abundant following FSH treatment (and JAK3 activation) as shown in figure 6d, while JNX treatment (and JAK3 inhibition) resulted in the reduction in CDKN1B phosphorylation.

In addition, we detected phosphorylation modifications for a specific region of CDKN1B, which is related to CDKN1B inactivation (Figure 8b; peptide 7 shown). In contrast to JAK3 phosphorylation, which activate the protein, phosphorylation of Threonine (T) residues at positions 187 and 198 of CDKN1B are an indication of CDKN1B inactivation, which results in cell cycle progression and cellular growth. As shown in Figure 6, CDKN1B total abundance decreased following FSH treatment, while JAK3 inhibition following JNX treatment resulted in an increase in total CDKN1B abundance. These data provide evidence that FSH stimulates JAK3 phosphorylation and activity and suppresses CDKN1B activity, which allows GC proliferation as demonstrated by the increase in proliferation markers.

## Discussion

The JAK/STAT signaling pathway begins with extracellular binding of members of a family of structurally related cytokines, interleukins, interferons, colony-stimulating factors, and some hormones to their corresponding structurally related transmembrane receptors as previously described^43,44^. This enables transactivation of receptor-bound JAKs that catalyze tyrosine phosphorylation of receptors and STATs, resulting in the formation of homodimers and/or heterodimers that accumulate in the nucleus and regulate gene transcription. JAK/STAT signaling pathways are major players in controlling cellular proliferation and differentiation processes. From this study, we show that FSH-induced phosphorylation of JAK3 (revealed via UHPLC-MS/MS) and overexpression of JAK3 seem to directly affect protein abundance and phosphorylation of downstream target STAT3. STAT proteins and in particular STAT3 are involved in transducing regulatory signals initiated following activation of JAK3 within the JAK/STAT signaling pathway to regulate the expression of specific genes associated with proliferation, migration and survival^27,45,46^. Upon activation of JAK3, the non-phosphorylated STAT3 proteins are recruited and phosphorylated, then dimerize and translocate into the nucleus where they bind to specific regions within the DNA and regulate expression of specific target genes. In contrast, inactivation of JAK3 protein in GC following JNX treatment significantly decreased STAT3 phosphorylation, which could alter the JAK/STAT signaling pathway in these cells. This is in accordance with previous studies showing downregulation of pSTAT3 following inhibition of JAK3^47^. The positive effect of FSH in JAK3 activation and STAT3 phosphorylation supports the activation of JAK/STAT pathway by FSH as well as the potential role of JAK3 activation in regulating GC proliferation, steroidogenic activity and survival through the JAK/STAT signaling and altering downstream targets activity. We have shown that FSH stimulates the expression of proliferation-associated genes *CCND2* and *PCNA* and differentiation-associated genes *CYP11A1* and, to a lesser degree, *CYP19A1,* which encode for enzymes involved in steroidogenic activity of granulosa cells^48–51^. These genes are involved in different stages of follicular development and growth and are associated with several intracellular signaling pathways including the JAK/STAT signaling pathway.

It has been reported that type II cytokine receptors are associated with JAK3 phosphorylation leading to the recruitment and phosphorylation of target proteins required for cell proliferation and survival^27,45^. JAK3 was shown to be involved in mediating signals initiated by cytokine signaling through coupling with the common γ chain of receptors for interleukins (IL)2, IL4, IL7, IL9, IL15 and IL21 and subsequently to play a critical role in the development, proliferation and differentiation of immune cells^44,52,53^. JAK3 is specifically responsive to these cytokines since it only binds to the common γ chain. Interleukins, which are key mediators of immune responses, may affect mechanisms crucial for granulosa cells function including their maturation and differentiation depending on the presence or not of FSH and thus may play a regulatory role in reproductive function^54^. It was shown that less differentiated granulosa cells from small follicles are more responsive to cytokines than are highly differentiated granulosa cells within large follicles^55^. Similarly, small follicles are also more responsive to FSH than large follicles while JAK3 was shown to be differentially expressed in small and growing dominant follicles^15,17^. FSH affects follicular growth, maturation, dominant follicle selection as well as estradiol production ^56^. On the other hand, growth factors and their receptors may affect the signaling network that is commonly activated by FSH receptors, which eventually may activate several signaling pathways including ERK1/2, the phosphatidylinositol-4,5-bisphosphate 3-kinases (PI3K)/protein kinase B (AKT)^57,58^ and also the JAK/STAT pathway as suggested in this study. UHPLC-MS/MS data showed that FSH induced JAK3 phosphorylation at specific residues, which are associated with JAK3 activation and related to increased cell proliferation as previously described^59,60^. These observations imply a synergically-induced effect of FSH and cytokines in granulosa cells that initiate a cascade of actions leading to JAK3 autophosphorylation and activation and recruitment of downstream target proteins.

A more accurate determination of the number and function of FSH-regulated genes as well as LH-regulated genes have been reported over the years^15,16,61-63^ and demonstrate the importance of functional studies during the later stages of follicular development to better coordinate the ovarian activity. In this regard, we have shown previously the suppression of JAK3 *in vivo* by endogenous LH or hCG injection in bovine species and identified JAK3 binding partners in GC using the yeast two-hybrid method^15,17^. We hypothesized then that, in contrast to LH downregulatory effects on JAK3 expression, FSH could induce JAK3 expression/activation, which in turn would regulate GC function by activating the JAK/STAT signaling pathway and modulate signaling pathways associated with JAK3 binding partners such as CDKN1B, which negatively affects cells proliferation if not phosphorylated, and MAPK8IP3 to maintain GC proliferation and follicular development. The data reported here support this hypothesis since inhibition of JAK3 led to a decrease in GC proliferation shown by a reduction in cell proliferation markers as well as disturbing the steroidogenic activity of GC. In contrast, JAK3 phosphorylation by FSH led to an increase in CDKN1B phosphorylation at specific target amino acids, while JAK3 inhibition resulted in a decrease in MAPK8IP3 abundance and phosphorylation.

CDKN1B (p27^Kip1^) is a cyclin-dependent kinase inhibitor that binds to and prevents the activation of cyclin E-CDK2 or cyclin D-CDK4 complexes, and thus controls the cell cycle progression at G1^64,65^. The degradation of CDKN1B, which is triggered by its CDK-dependent phosphorylation and subsequent ubiquitination, is required for the cellular transition from the quiescence to the proliferative state^66^. Therefore, CDKN1B acts either as an inhibitor or an activator of cyclin type D-CDK4 complexes depending on its phosphorylation state and/or stoichiometry. The phosphorylation of CDKN1B occurs on serine, threonine and tyrosine residues. Phosphorylation on S^10^ is the major site of phosphorylation in resting cells and takes place at the G(0)-G(1) phase leading to protein stability^67^. Phosphorylation on other sites is potentiated by mitogens or growth factors and in certain cancer cell lines^68,69^, meaning that phosphorylated CDKN1B found in the cytoplasm is inactive. Consistent with these observations, we showed in this study CDKN1B phosphorylation following FSH treatment and subsequent JAK3 activation. This observation is concomitant with increased JAK3 total abundance and phosphorylation level suggesting that CDKN1B phosphorylation at specific residues is necessary following FSH addition in order to allow progression of the cell cycle and GC proliferation. Based on these reports, and in light of our current findings, it may be possible that CDKN1B binding to JAK3 leads to its phosphorylation and inactivation in granulosa cells of small and growing dominant follicles, thus allowing follicular development.

Additionally, we previously reported the strongest expression of CDKN1B in the corpus luteum^17^ suggesting a role for CDKN1B in establishing the non-proliferative state, which is required for differentiation or for proper functioning of the differentiated luteal. Our findings are therefore consistent with previous results showing that the luteinization process is associated with up-regulation of *CDKN1B* that accumulated during initial phases of luteinization and remained elevated until termination of the luteal function^70^. It was also demonstrated that inhibition of JAK3/STAT3 signaling significantly decreased viability of colon cancer cells due to apoptosis and cell-cycle arrest through down-regulation of cell cycle genes including cyclin D2 and up-regulation of CDKN1B^47^. Our data show similar findings following JAK3 inhibition with decreased expression of cell proliferation markers *cyclin D2* and *PCNA,* decreased STAT3 phosphorylation and increased CDKN1B protein abundance, confirming the biological importance of JAK3 and the JAK/STAT pathway in the ovarian function. Moreover, our findings confirmed that downstream targets of the JAK/STAT signaling that are involved in cell-cycle regulation, including cyclin D2 and CDKN1B, also play a major role in the regulation of GC proliferation.

Conversely, MAPK8IP3 (JNK-interacting protein 3; JIP3) is a mitogen activated tyrosine kinase (MAPK) that contributes to the C-Jun signalling pathway. The different classes of MAPKs play important roles in various cellular processes including cell proliferation, differentiation and apoptosis. JNKs are activated by diverse stimuli including DNA damage or inflammatory cytokines^71^. MAPK activation may be facilitated by the formation of signaling modules, and it has been well established that scaffold proteins play important roles by interacting with MAPKs and their upstream kinases^72^. MAPK8IP3 has been reported as a scaffold protein within the JNK signaling cascade^73^. We show here that MAPK8IP3 exhibited increased phosphorylation of residues associated with cell proliferation. Although FSH did not increase the overall total MAPK8IP3 abundance, JAK3 inhibition by JNX resulted in a notable decrease in total abundance of MAPK8IP3. This insight might show a direct effect of JAK3 inhibition on the activation of MAPK8IP3, therefore affecting other associated signaling pathways in GC.

The data from this study show that FSH regulates expression of genes involved in GC proliferation and follicular development. Most importantly, these findings implicate FSH in modulating JAK3 phosphorylation and the JAK/STAT signaling pathway likely through phosphorylation of STAT3 proteins. The results also demonstrate a crucial role of JAK3 in GC proliferation and modulation of downstream targets CDKN1B and MAPK8IP3 in response to a hormonal signal (FSH) leading to follicular growth and development. Although the exact mechanisms by which FSH affects JAK3 and regulates STAT3 and the downstream targets were not fully elucidated, this study serves as a basis for studies targeting GC regulation during late stages of follicular development and in JAK3-inhibited GC to identify pathways and downstream targets affected by FSH-induced genes and more specifically JAK3.

## Data availability statement

All data pertinent to this study are contained in the article or available upon request. For all requests, please contact Dr. Kalidou Ndiaye.

## Acknowledgements

This study was supported by a Discovery Grant from the Natural Sciences and Engineering Research Council of Canada (NSERC) (RGPIN#04516) to K. Ndiaye. Proteomics equipment was funded by the Canadian Foundation for Innovation (CFI) and the Fonds de Recherche du Québec (FRQ), the Government of Quebec (F. Beaudry CFI John R. Evans Leaders grant no. 36706). The funder had no role in study design, data collection, and analysis, decision to publish, or preparation of the manuscript.

The authors thank Dr. Jacques G. Lussier for all the advice, suggestions, and critical comments on successful progress of this project.

## Author contributions

Investigation, A.Z. and K.N.; Visualization, A.Z. and K.N.; Validation, A.Z. and K.N.; Formal analysis, A.Z. and K.N.; Proteomics analyses, F.B., A.Z. and K.N.; Writing-Original draft preparation, A.Z.; Methodology, A.Z., F.B. and K.N.; Conceptualization, K.N.; Resources, K.N.; Writing-Review and Editing, A.Z., F.B. and K.N.; Supervision, K.N.; Project Administration, K.N.; Funding Acquisition, K.N and F.B. All authors have read and agreed to the published version of the manuscript.

## Additional information

### Competing interest

The authors declare that there is no conflict of interests.

## Notes

### Competing Interest Statement

The authors have declared no competing interest.

